# TGF-β signaling suppresses TCA cycle metabolism in renal cancer

**DOI:** 10.1101/2021.02.19.429599

**Authors:** Hyeyoung Nam, Anirban Kundu, Suman Karki, Garrett Brinkley, Darshan S. Chandrashekar, Richard L. Kirkman, Juan Liu, Maria V. Liberti, Jason W. Locasale, Tanecia Mitchell, Sooryanarayana Varambally, Sunil Sudarshan

**Affiliations:** Departments of Urology, University of Alabama at Birmingham, Birmingham, AL, 35294; Pathology University of Alabama at Birmingham, Birmingham, AL, 35294; Department of Pharmacology and Cancer Biology, Duke University, Durham, NC 27710; Birmingham Veterans Affairs Medical Center, Birmingham, AL, 35233

## Abstract

The Warburg effect is one of most-well studied metabolic phenomenon in cancer cells. For the most part, these studies have focused on enhanced rates of glycolysis observed in various models. The presumption has been that mitochondrial metabolism is suppressed. However, recent studies indicate that the extent of mitochondrial metabolism is far more heterogeneous in tumors than originally presumed. One tumor type with suppression of mitochondrial metabolism is renal cell carcinoma (RCC). Prior studies indicate that suppressed TCA cycle enzyme mRNA expression is associated with aggressive RCC. Yet, the mechanisms that regulate the TCA cycle in RCC remain uncharacterized. Here, we demonstrate that loss of TCA cycle enzyme expression is retained in RCC metastatic tissues. Moreover, proteomic analysis demonstrates that reduced TCA cycle enzyme expression is far more pronounced in RCC relative to other tumor types. Loss of TCA cycle enzyme expression is correlated with reduced expression of the transcription factor peroxisome proliferator-activated receptor gamma coactivator 1-alpha (PGC-1α) which is also lost in RCC tissues. PGC-1α re-expression in RCC cells restores the expression of TCA cycle enzymes *in vitro* and *in vivo* and leads to enhanced glucose carbon incorporation into TCA cycle intermediates. Mechanistically, TGF-β signaling, in concert with histone deacetylase 7 (HDAC7), suppresses TCA cycle enzyme expression. In turn, pharmacologic inhibition of TGF-β restores expression of TCA cycle enzyme expression and suppresses tumor growth in an orthotopic model of RCC. Taken together, our findings reveal a novel role for the TGF-β /HDAC7 axis in global suppression of TCA cycle enzymes in RCC and provide novel insight into the molecular basis of altered mitochondrial metabolism in this malignancy.

## INTRODUCTION

Clear cell renal cell carcinoma (ccRCC) is the most common histologic subtype of kidney cancer. Approximately 30% to 40% of patients with ccRCC present with metastases at initial diagnosis [1, 2]. Individuals with organ-confined RCC tumors are considered to have an excellent prognosis with treatment. In contrast, patients with advanced disease have poor survival rates. Thus, a better understanding of the factors leading to tumor progression in RCC and the development of novel therapeutic strategies are of potential significance. ccRCC is known to have striking metabolic features. The most-well characterized is increased expression of glycolytic genes that results from loss/mutation of *VHL*, a common tumor initiating event. Loss of the E3 ubiquitin ligase activity of VHL results in stabilization of the hypoxia inducible factors (HIFs) and subsequent upregulation of hypoxia-responsive genes [3–5]. Glycolytic enzymes are known HIF transcriptional target in cancer [6, 7].

While enhanced glycolysis is a shared feature of many caner types, the expression of enzymes involved in mitochondrial metabolism is more heterogeneous. Notably, ccRCC is among the tumor types with the most prominent decrease in the mRNA expression of TCA cycle enzymes [8]. In agreement, we previously noted reduced expression of fumarate hydratase (FH) protein in ccRCC [9]. Moreover, a recent stable isotope labelling study in ccRCC patients demonstrated reduced incorporation of glucose-derived carbons into TCA cycle metabolites[10]. These data are among the most compelling to demonstrate reduced TCA cycle metabolism in kidney cancer. The decreased mRNA expression of TCA cycle enzymes in kidney tumor likely has biological relevance. A major conclusion of the TCGA (The Cancer Genome Atlas) analysis of kidney cancer was a metabolic shift in aggressive tumors marked by the down-regulation of genes encoding enzymes of the TCA cycle including *FH* (fumarate hydratase), *ACO2* (aconitase), *SUCLG1* (succinate-CoA ligase, α subunit), *OGDH* (oxoglutarate dehydrogenase)[11]. Despite these findings, the molecular basis by which the TCA cycle is altered in ccRCC remains poorly understood. As a result, the implications of this alteration as it pertains to RCC tumor biology remains unknown.

We recently reported an integrative analysis on RCC tumor progression which included normal kidney, primary tumors as well as metastatic tissues [12]. This analysis demonstrated that loss of mRNA expression of *PPARGC1A*, which encodes for the transcription factor PGC-1α, as among the most suppressed genes. PGC-1α was originally identified as a transcriptional coactivator involved in mitochondrial function and thermogenesis in brown fat [13]. *PPARGC1A* is known to be expressed in metabolically active tissues such as the kidney. Recently, we identified a novel role for PGC-1α in suppressing both the expression of collagen genes and tumor progression in an orthotopic model of RCC[14]. Prior studies have demonstrated that the transcriptional regulation of collagen gene expression is mediated by transforming growth factor beta (TGF-β) [15]. Although TGF-β has been implicated in invasive behaviors in many cancers (including RCC) via its promotion of the epithelial to mesenchymal transition (EMT), its role in mitochondrial TCA cycle metabolism is undefined. These data led us to consider the interrelationship between TGF-β and PGC-1α in the context of RCC and the relevance of this axis to RCC metabolism.

Here, we demonstrate that global repression of TCA cycle enzymes is a unique feature of RCC which is also found in RCC metastatic deposits. Mechanistically, TGF-β and histone deacetylase 7 (HDAC7) cooperate to repress PGC-1α and TCA cycle enzyme expression. Moreover, pharmacologic inhibition of TGF-β can restore TCA cycle enzyme expression *in vivo*. Overall, our findings provide novel insight into the epigenetic basis of altered mitochondrial metabolism in RCC. Moreover, they are among the first data to demonstrate that mitochondrial aspects of classic Warburg metabolism are pharmacologically targetable in RCC.

## RESULTS

### The expression of TCA cycle enzymes is lost in ccRCC

Prior TCGA analyses demonstrated reduced mRNA expression of genes encoding TCA cycle enzymes. Apart from our previous study on fumarate hydratase (FH) [9], these data have not been validated at the protein level. We first examined primary ccRCC specimens and patient-matched adjacent normal kidney and found reduced protein levels of aconitase 2 (ACO2) and the alpha subunit of succinate-CoA ligase (SUCLG1) in RCC (Fig. 1A). In addition, we observed the relative expression of TCA cycle enzymes in a panel of RCC cell lines. RCC cell lines had decreased protein levels of ACO2 and SUCLG1 relative to RPTEC renal proximal tubule epithelial cells (Fig. 1B). We analyzed publicly available proteomics data released from TCGA (CPTAC) using the UALCAN analysis portal [16]. These data validated our findings of reduced protein expression of TCA cycle enzymes in ccRCC relative to normal kidney. (Fig. 1C). Analysis of proteomics data from other tumor types demonstrated that reduced expression of TCA cycle enzymes is far more pronounced in ccRCC relative to other tumor types (Fig.1C and S1). Whereas breast and colon cancers demonstrated slightly reduced expression, no changes were noted in ovarian and uterine cancers. *VHL* alterations are the most common tumor initiating event in ccRCC. An established sequelae of *VHL* loss is stabilization of hypoxia inducible factors (HIFs) which has been linked with alterations in mitochondrial metabolism [17]. However, we noted that both *VHL* null (RCC4 and RCC10) as well as *VHL* WT lines (CAKI-1 and RXF-393) had reduced expression of TCA cycle enzymes (Fig. 1B). We therefore assessed the relative mRNA expression of TCA cycle enzymes by *VHL* status in ccRCC in the TCGA data set (Fig. 1D). Consistent with our findings in cell lines, we found that both *VHL* WT and mutant tumors demonstrated reduced mRNA expression of TCA cycle enzymes relative to normal kidney (Fig. 1D). We next considered whether these changes were maintained in RCC metastatic tissues as TCGA only analyzed primary tumors. We recently reported the gene expression landscape of ccRCC progression which encompassed transcriptomic analysis of normal kidney (n = 9), primary RCC (n = 9), and metastatic RCC tissue deposits (n = 26) [12]. Notably, we find that the mRNA expression of TCA cycle enzymes is reduced in metastatic tissues indicating that the pronounced shift in TCA cycle metabolism is retained with tumor progression (Fig. 1E). As these samples were not patient-matched, we examined the expression of TCA cycle enzymes in a separate cohort which included patient-matched samples of normal kidney, primary tumor, and metastatic tissue. This analysis confirmed that the reduced mRNA expression of TCA cycle enzymes (i.e. *ACO2*, *OGDH*, *SUCLG1*, and *MDH2*) in primary tumor was retained in metastatic tissues (Fig. 1F).

**Figure 1.**
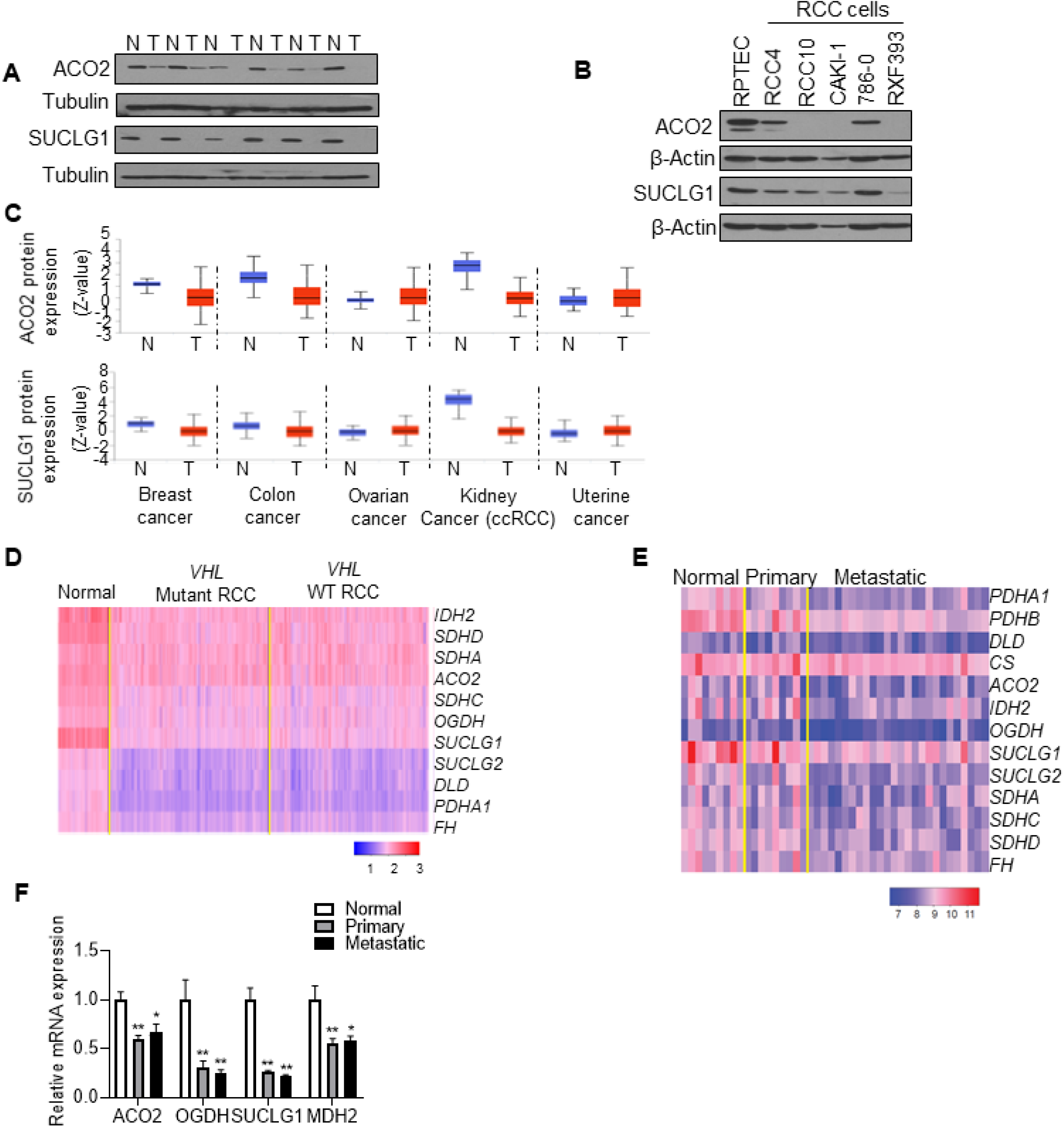
Suppression of TCA cycle enzymes in ccRCC. **A)** lmmunoblot analysis of ACO2 and SUCLG1 in patient­ matched normal kidney (N) and tumor (T). **B)** Western blot analysis of ACO2 and SUCLG1 in a panel of RCC cell lines relative to RPTEC primary renal proximal tubule epithelial cells. **C)** Protein expression of ACO2 and SUCLG1 across pan-cancer subtypes from the CPTAC cohorts. Data were analyzed with the UALCAN analysis portal. **D)** Heatmap representing expression patterns of TCA cycle enzymes in normal kidney (n=72), *VHL* mutant RCC (n=224), and *VHL* WTRCC (n=225). Data were extracted fromTCGA KIRC dataset.The heatmap shows the log10-transformedTPM values for each gene. **E)** Heatmap representing expression patterns ofTCA cycle enzymes using the lllumina Human HT-12 v4 bead array in the three-patient groups (normal n=9, primary n=9, and metastasis n=26). Colors in the heatmap represent log transformed quantile normalized expression values for selected set of probes in individual samples. **F)** Relative mRNA expression ofTCA cycle enzymes in a separate cohort of patient-matched samples. Transcript levels were normalized to those of *TBP* (n=6/group). Asterisks indicate significant differences compared to normal kidney (\**P*<0.05, \*\**P*<0.01, One-way ANOVA with Tukey’s multiple comparisons test).

### The transcriptional landscape of ccRCC reveals positive correlation between transcripts encoding TCA cycle enzymes and *PPARGC1A*

We next wanted to gain insight into the mechanism that drive the suppression of TCA cycle enzyme in RCC. We recently reported that *PPARGC1A* expression is progressively silenced with RCC tumor progression [14]. *PPARGC1A* encodes for the transcription factor peroxisome proliferator-activated receptor-gamma coactivator 1-alpha (PGC-1α). Restoration of PGC-1α suppresses *in vivo* tumor progression in an orthotopic model of RCC [14]. PGC-1α is known to have a role in mitochondrial metabolism. We first performed correlation analysis of genes positively correlated with *PPARGC1A* in metastatic RCC tissues. KEGG analysis demonstrated that metabolic pathways including TCA cycle enzymes were among the most enriched pathways (Fig. 2A). Several genes of the TCA cycle enzyme demonstrated a statistically significant positive correlation with *PPARGC1A* including *ACO2*, *SUCLG1*, *SDHA* (succinate dehydrogenase A), *SDHB* (succinate dehydrogenase B), *SUCLA2* (succinate-CoA ligase ADP-forming subunit beta), and *DLAT* (dihydrolipoamide S-acetyltransferase) (Fig. 2B and 2C). Furthermore, a positive correlation between *PPARGC1A* and several TCA cycle enzyme genes was also observed upon analysis of TCGA data (Fig. 2D and 2E).

**Figure 2.**
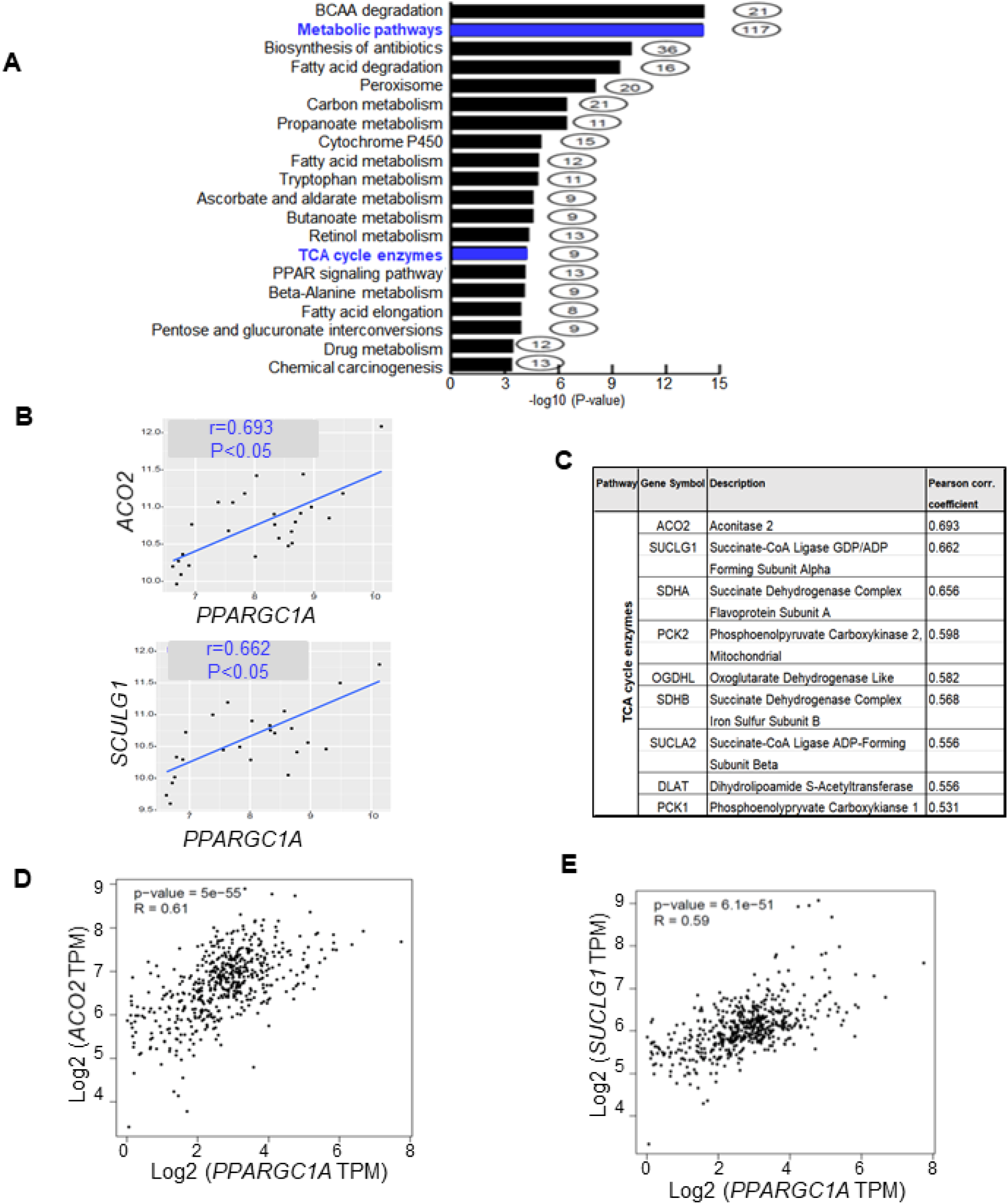
*PPARGC1A* expression is positively correlated with the expression of TCA cycle enzymes in ccRCC. **A)** KEGG pathway analysis represent the pathways that are positively correlated with *PPARGC1A* in metastatic RCC samples (n=26). B) Positive correlation between *PPARGC1A* and TCA cycle enzymes in metastatic tumor samples determined by Pearson’s correlation (n=26, *P*<O.O5). C) Positive correlation between *PPARGC1A* and several TCA cycle enzyme genes. **D and E)** Results of correlation analysis between *PPARGC1A* and TCA cycle enzymes (*AC02* and *SUCLG1*) in RCC samples from TCGA KIRC dataset. Data extracted using GEPIA web server.

### *PPARGC1A* re-expression restores the expression of TCA cycle enzymes and mitochondrial function

The correlation studies led us to consider the role of PGC-1α loss on the expression of TCA cycle enzymes. Consistent with the decreased mRNA expression of *PPARGC1A*, RCC cell lines demonstrated a significant decrease in mRNA expression of TCA cycle enzymes including *ACO2*, *OGDH*, and *SUCLG1* relative to normal kidney (Fig. 3A). PGC-1α expression was restored in CAKI-1 and RXF393 RCC cells. PGC-1α restoration led to increased protein expression of ACO2 and SUCLG1 in RCC cells (Fig. 3B). Consistent with the protein data, PGC-1α re-expression significantly increased the mRNA expression of TCA cycle enzymes in both RCC cells (Fig. 3C). Similar findings were found in RCC4 cells transduced with adenovirus containing cDNA for PGC-1α (Fig. S2A and S2B). Knockdown of *PPARGC1A* in RCC4 cells, which have a low but detectable level of PGC-1α [14], led to reduced TCA cycle enzyme expression (Fig. 3D). Moreover, these findings were relevant *in vivo* as restoration of PGC-1α in SN12PM6-1 RCC cells led to increased mRNA and protein levels of TCA cycle enzymes in orthotopic xenografts (Fig. 3E and 3F).

**Figure 3.**
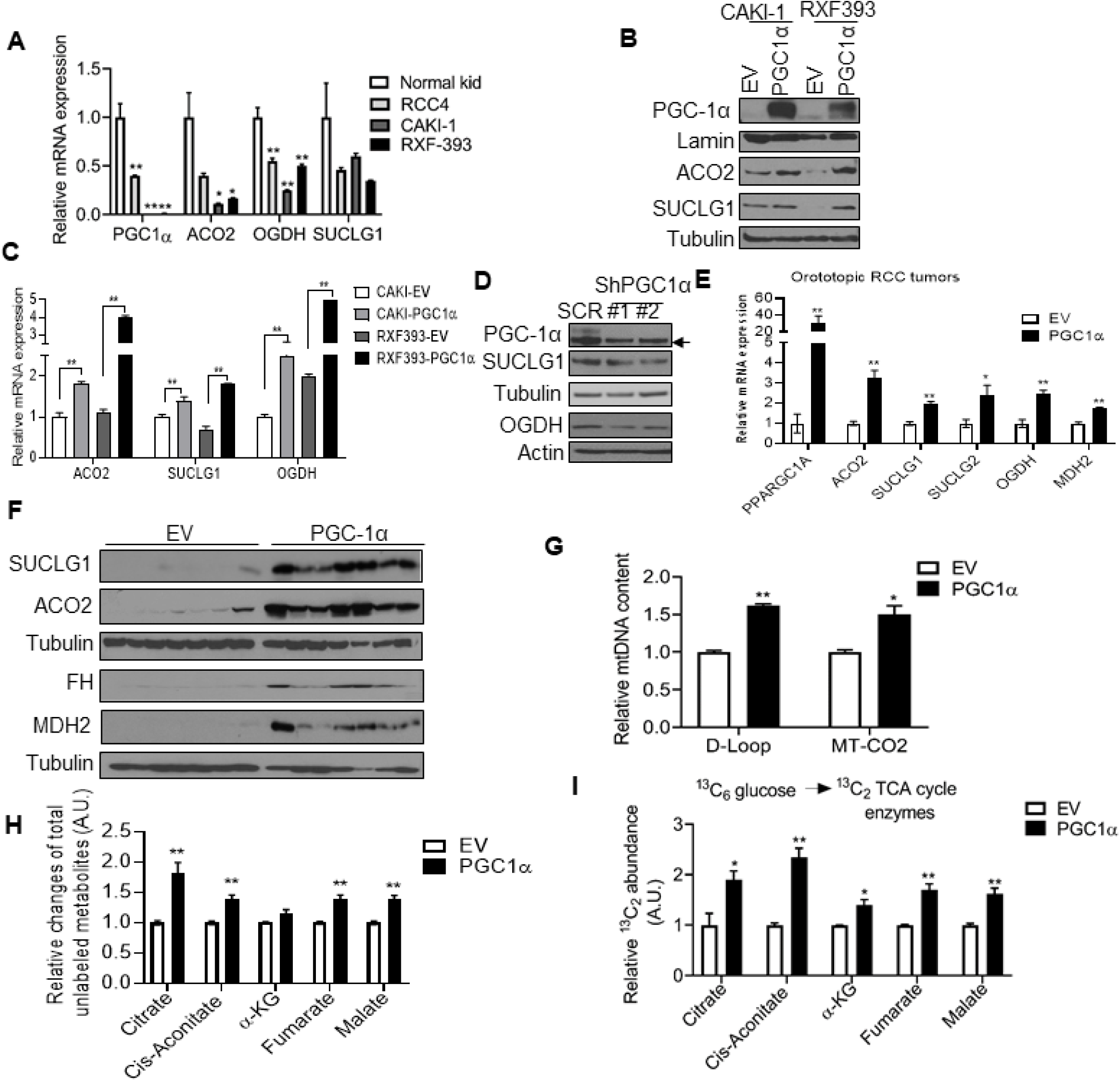
PGC-1 a re-expression up-regulates the expression of TCA cycle enzymes and glucose flux to TCA cycle enzymes. **A)** Relative mRNA expression of the indicated genes in a panel of RCC cells relative to normal kidney (n=3/group). Transcript levels were normalized to those of *TBP.* **B)** Western blot analysis of the indicated proteins in RCC cells stably expressing either EV or PGC-1a. C) Representative mRNA expression of TCA cycle enzymes from RCC cells transduced either EV or PGC1a (n=3/group). All data represent 3 independent experiments and error bars are SEM. Asterisks indicate significant differences compared to control (\**P*<0.05,\*\**P*<0.01,One-way ANOVA with Tukey’s multiple comparisons test or 2 tailed student *I* test). **D)** Western blot analysis of the indicated proteins in RCC4 cells stably expressing shRNA control (SCR) or two independent PGC-1a shRNA constructs. Arrow represents non-specific band. **E and F)** SN12PM6-1 cells stably expressing either EV or PGC1 a were orthotopically implanted into the left kidney of SCIO mice. At 6 weeks from tumor challenge, kidney tissues were harvested and analyzed for the expression of **E)** mRNA and **F)** protein for the indicated genes (n=7/group). **G)** CAKl-1 cells stably expressing EV or PGC-1α were measured for the amount of mtDNA D-Loop structure and *MT-CO2* gene, encoding for mtDNA-encoded cytochrome C oxidase II (MT-CO2). qPCR was performed with total DNA. **H and I)** CAKl-1 cells stably expressing with EV or PGC-1a were incubated with uniformly labeled [U-^13^C_6_] glucose for 24 h. The relative levels of **H)** total unlabeled metabolites and **I)** labeled TCA cycle intermediates (M+2) were analyzed using LC-MS (n=3/group).

We next examined the effect of PGC-1α on mitochondrial DNA content. The amount of both mitochondrial DNA D-Loop structure and *MT-CO2* gene, encoding for mtDNA-encoded cytochrome C oxidase II (MT-CO2), was significantly increased in PGC-1α expressing RCC cells (Fig. 3G). We next assessed the effects of PGC-1α on TCA cycle activity with LC-MS analysis using uniformly labeled [U-^13^C_6_] glucose. First, we noted that PGC-1α increased total unlabeled pools of multiple TCA cycle metabolites including citrate, cis-aconitate, fumarate, and malate (Fig. 3H). Furthermore, PGC-1α led to increased labeling of TCA cycle metabolites indicating the enhanced contribution of glucose-derived carbons to the TCA cycle (Fig. 3I). Collectively, these data indicate that PGC-1α restoration in RCC cells can promote the expression of TCA cycle enzymes as well as increase mitochondrial DNA content and enzyme activity.

### Blockade of TGF-β signaling rescues the expression of *PPARGC1A* and mitochondrial function in RCC

The profound role of PGC-1α on the expression of TCA cycle enzymes led us to consider the mechanism that promotes loss of PGC-1α expression in RCC. Prior studies indicated that loss of PGC-1α is HIF-dependent in RCC [18]. However, our analysis of TCGA data demonstrates that TCA cycle enzyme expression is reduced irrespective of *VHL* status indicating an alternate mechanism (Fig. 1D). We recently reported the increased mRNA expression of collagen (*COL*) family members is highly associated with RCC metastasis and that PGC-1α restoration suppresses the expression of *COLs* [14]. Prior studies have indicated a role for transforming growth factor beta (TGF-β) in promoting *COL* gene expression as part of the EMT program [15]. We therefore assessed TGF-β’s role in regulating the expression of PGC-1α. Consistent with TGF-β’s role in promoting *COL* expression, we find that pharmacologic inhibition of TGF-β suppresses the mRNA expression of several *COL* genes including *COL1A1*, *COL5A1*, *COL5A2*, and *COL11A1* in CAKI-1 and RXF-393 cells (Fig. S3A and S3B). Consistent with the mRNA data, inhibition of TGF-β led to decreased protein expression of COL1A1 in RCC cells (Fig. S3C). Inhibition of TGF-β signaling utilizing multiple inhibitors (SB431542, LY2109761, and LY364947) resulted in an increase of *PPARGC1A* transcript levels in RCC cells (Fig. 4A and S3D). In addition, TGF-β inhibition led to a significant increase in mRNA expression of the TCA cycle enzymes in 769-P (Fig. 4B) and CAKI-1 cells (Fig. 4C). Accordingly, increased protein levels of ACO2 and SUCLG1 were observed in CAKI-1 cells treated with TGF-β inhibitors (Fig. 4D). In addition, HK2 renal epithelial cells treated with TGF-β led to reduced protein expression of TCA cycle enzymes (Fig. 4E). Based on these observations, we examined the impact of TGF-β inhibition on cellular bioenergetics. We measured the oxygen consumption rate (OCR) of RCC cells treated with either control (DMSO) or two pharmacological TGF-β inhibitors (Fig. 4F). Notably, both TGF-β inhibitors increased cellular bioenergetics in RCC cells compared to control cells (Fig. 4F). In particular, both inhibitors significantly increased basal respiration, ATP-linked respiration (assessed following oligomycin treatment), and maximal respiration (assessed following treatment with uncoupler FCCP) (Fig. 4G). SB431542 significantly increased non-mitochondrial respiration (following antimycin A treatment), whereas LY364947 had no effect (Fig. 4G). Collectively, these data demonstrate a role for TGF-β signaling in regulating both TCA cycle enzyme expression and cellular bioenergetics in RCC cells.

**Figure 4.**
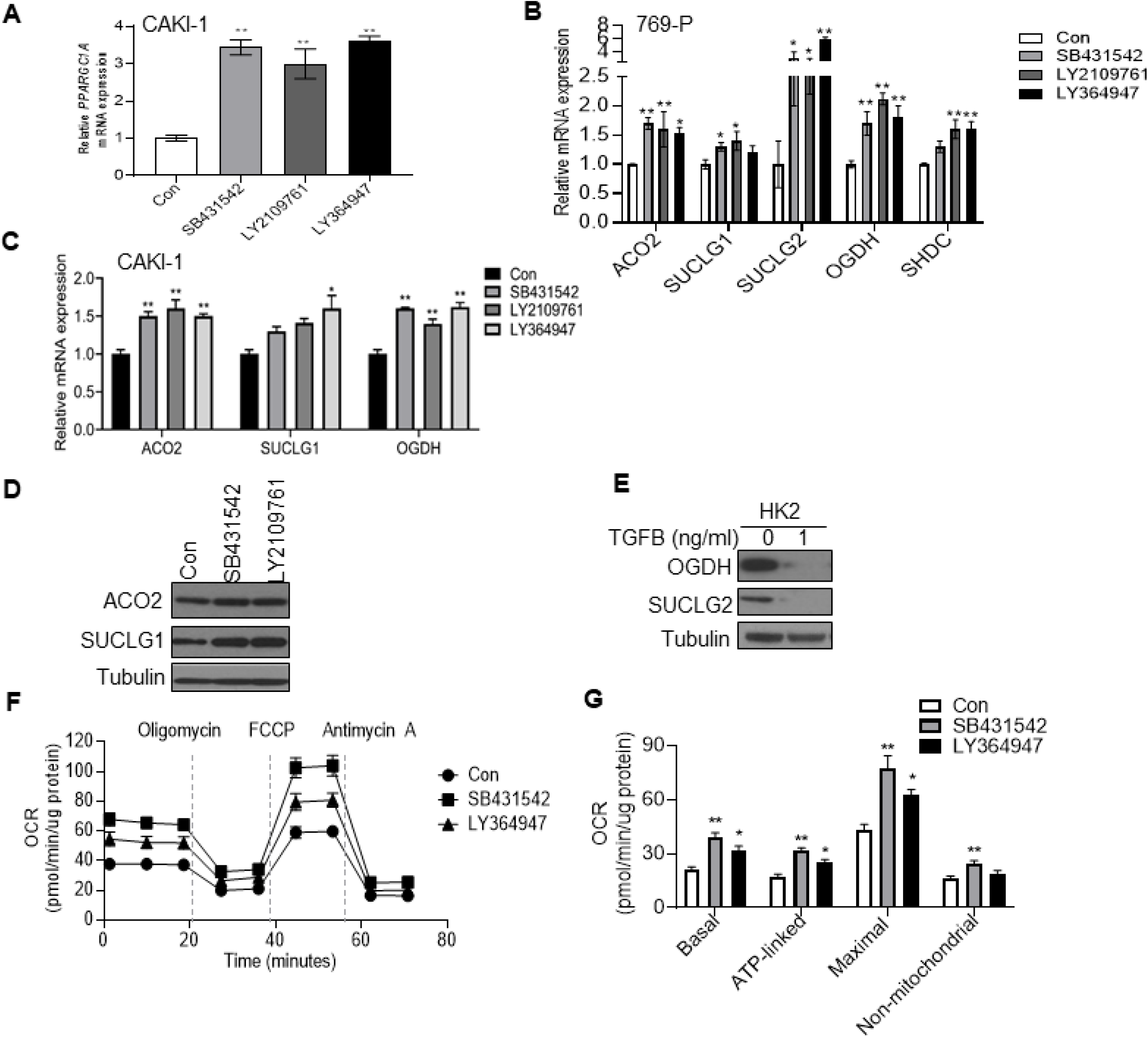
The expressions of *PPARGC1A* and TCA cycle enzymes are restored by blockade of TGFB signaling. **A)** Relative mRNA expression of *PPARGC1A* in CAKl-1 cells treated with either DMSO (Con) or indicated pharmacological TGF-13 inhibitors (10 μM) for 48 h (n=3/group). **Band C)** Relative mRNA expression of TCA cycle enzymes in 769-P and CAKl-1 cells treated with either DMSO or indicated TGF-13 inhibitors (10 μM) for 48 h (n=3/group). **D)** lmmunoblot analysis for ACO2 and SUCLG1 protein expression in cell lysates from CAKl-1 cells treated with indicated TGF-13 inhibitors for 48 h. **E)** HK2 cells, an immortalized proximal tubule epithelial cell line, were exposed to TGFB (1 ng/ml)for 24 h and followed by immunoblot analysis for OGDH and SUCLG2 expression. **F and G)** Oxygen consumption rate (OCR) was analyzed in CAKl-1 cells treated with either DMSO (Con) or indicated pharmacological TGFB inhibitors (10 μM) for 48 h (n=8/group). Oligomycin (1.5 μg/ml), FCCP (0.6 μM), and Antimycin A (10 μM) were sequentially added to the cells. Representative **F)** cellular bioenergetic profiles and **G)** individual parameters are shown. All data represent 2-3 independent experiments and are presented as ± SEM. Asterisks indicate significant differences compared to control (\**P*<0.05,\*\**P*<0.01, One-way ANOVA with Tukey’s multiple comparisons test).

### TGF-β inhibition reverses metabolic phenotypes of RCC *in vivo*

As noted previously, a major finding from the TCGA analysis of ccRCC was that loss of TCA cycle enzyme expression is associated with aggressive tumors. Given our *in vitro* findings, we next assessed if these findings are relevant *in vivo* to assess if mitochondrial aspects of Warburg metabolism could be reversed by pharmacologic means. CAKI-1 luciferase expressing RCC cells were injected orthotopically into the renal subcapsular region of SCID mice. RCC tumor bearing mice were then intraperitoneally treated with either 20% DMSO (control) or TGF-β inhibitor SB431542 (10 mg/kg in 20% DMSO). After 5 weeks of treatment, mice treated with SB431542 demonstrated significantly lower tumor burden as demonstrated by both bioluminescent imaging (Fig. 5A) and tumor weight (Fig. 5B). Analysis of tumor explants demonstrated increased expression of *PPARCG1A* mRNA and PGC-1α protein in mice treated with SB431542 (Fig. 5C and 5D). Moreover, TGF-β inhibitor treated tumors demonstrated increased expression of TCA cycle enzymes at both the mRNA and protein levels (Fig. 5E and 5F). Collectively, these data demonstrate the role of TGF-β in regulating the TCA cycle in RCC and that this can be targeted by pharmacologic means *in vivo*.

**Figure 5.**
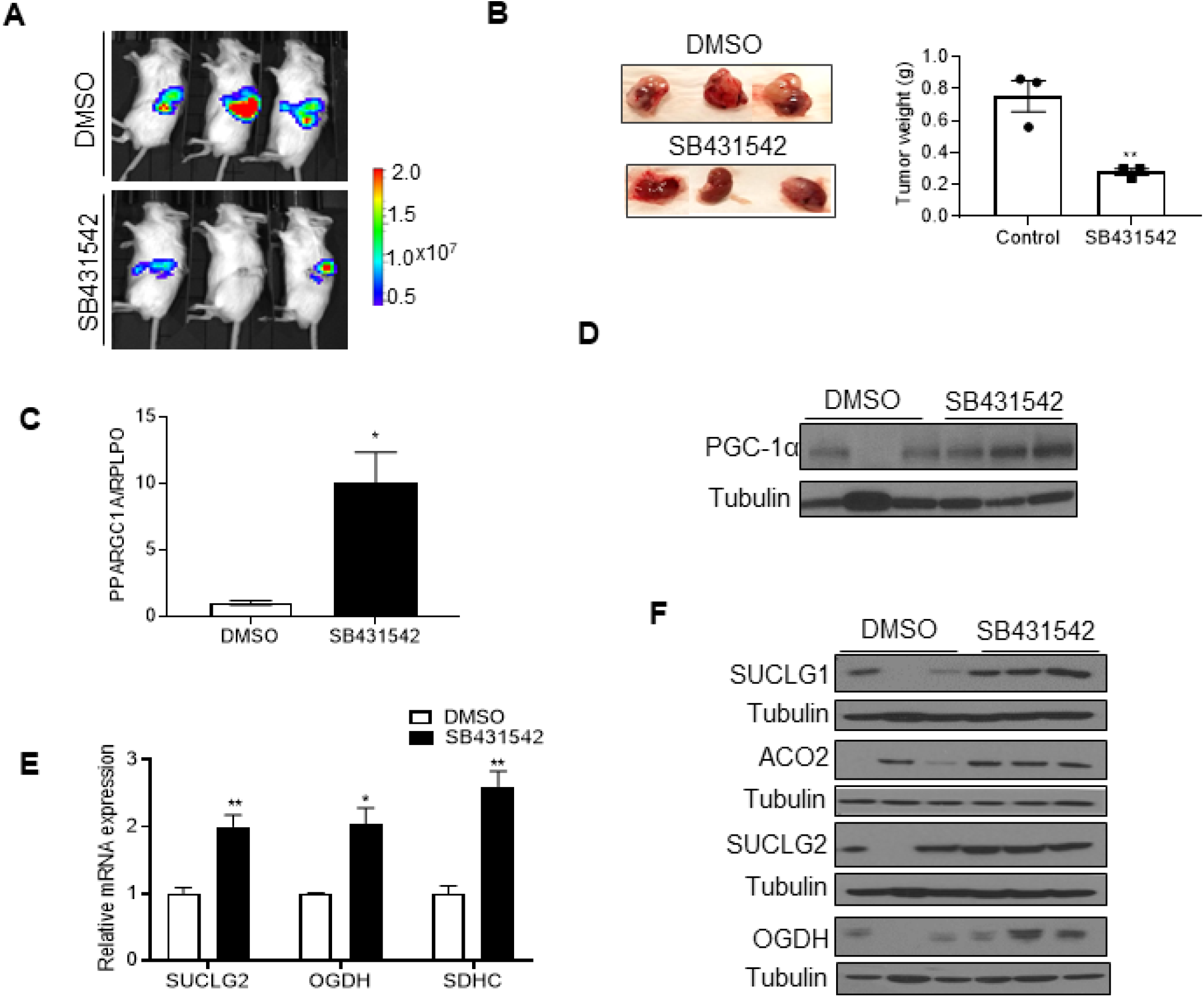
Inhibition of TGFB signaling suppresses tumor formation and restores the expression of *PPARGC1A* and TCA cycle enzymes *in vivo.* **A)** CAKl-1 luciferase expressing cells (1.5 × 10^6^) were implanted into renal subcapsular region of SCIO mice (n=3/group). The luciferase bioluminescence was used to monitor tumor growth. After confirming the presence of tumors one week after cell injection, the treatment with either DMSO or S8431542 (10 mg/kg in 20% DMSO) was intraperitoneally administrated three times per week for 5 weeks. The *in vivo* luciferase bioluminescence was taken at 5 weeks after treatment with either DMSO or S8431542. **B)** At 5 weeks after treatment, kidney tissues were harvested and weighted. **C)** The RNA was isolated from k dney tumors of mice treated with either DMSO or S8431542 and analyzed for mRNA expression of *PPARGC1A.* Transcript levels were normalized to those of *RPLPO.* **D)** lmmunoblot analysis of PGC1a in kidney tissues from the mice treated wi ith either DMSO or S8431542. **E)** Relative mRNA and **F)** protein expression of TCA cycle enzymes in kidney t ssues from the mice treated with either DMSO or S8431542. Transcript levels were normalized to those of *RPLPO.* All data are presented as± SEM (\**P*<0.05, \**P*<0.01, 2 tail student *t* test).

### HDAC7 acts as a corepressor for TGF-β mediated suppression of TCA cycle enzymes in RCC

We next investigated the mechanism by which TGF-β signaling repressed the expression of *PPARGC1A* and TCA cycle enzymes. TGF-β signaling regulates transcription by a complex network including SMAD proteins. SMAD complexes can inhibit transcription through interacting with transcriptional corepressors. Three SMAD corepressors have been identified including the homeodomain protein TG-interacting factors (TGIFs), Ski, and SnoN protein [19–21]. We therefore examined genes that are negatively correlated with these corepressors in the TCGA data set on ccRCC [22]. Intriguingly, this analysis identified that *TGIF2* is inversely correlated with the expression of genes involved in the TCA cycle enzymes. In fact, the TCA cycle was the top-ranked gene set (Tab. S3A). Based on these findings, we next performed loss of function studies via siRNA mediated knockdown of *TGIF2* in CAKI-1 cells. Knockdown of *TGIF2* led to significantly increased mRNA levels of TCA cycle enzymes including *ACO2*, *SUCLG1*, *OGDH*, and *SDHC* (Fig. 6A). Correspondingly, the protein levels of SUCLG1 were upregulated by knockdown of *TGIF2* (Fig. 6B). TGIF2 had been previously shown to be a transcriptional repressor by interacting with histone deacetylases (HDACs) including HDAC1 [21, 23]. We therefore assessed whether HDACs could contribute to silencing the expression of *PPARGC1A* and/or TCA cycle enzyme genes. We initially investigated the effects of the pan-HDAC inhibitor trichostatin A (TSA) on the expression of *PPARGC1A* in RCC cells. We found a significant increase in mRNA and protein expression of PGC-1α in RCC cells following treatment with TSA (Fig. 6C and 6D). Furthermore, TSA treatment increased the mRNA expression of TCA cycle enzymes in CAKI-1 cells (Fig. 6E). In concert, TSA treatment resulted in the increased protein expression of ACO2 and SUCLG1 in 769-P and CAKI-1 cells (Fig. 6F). We therefore assessed whether HDACs inversely correlated with TCA cycle enzymes (Tab. S3B). We found that *HDAC1* and *7* are negatively correlated with the mRNA expression of TCA cycle enzymes in the TCGA data set. We thus knocked down both HDAC1 and HDAC7. We confirmed target gene knockdown via RT-qPCR (Fig. S4A). We found that HDAC1 knockdown significantly induced the mRNA expression of *PPARGC1A* in 769-P cells (Fig. 6G). HDAC7 knockdown had a modest effect on *PPARGC1A* mRNA. However, HDAC7 knockdown led to a prominent increase in TCA cycle enzyme mRNA levels of *ACO2*, *SUCLG1*, and *SDHC* (Fig. 6G). In agreement with the mRNA data, HDAC7 knockdown demonstrated increased ACO2 and SUCLG1 protein expression in RCC (Fig. 6H). Furthermore, we investigated potential SMAD binding sites within 1 kb of the *SUCLG1* transcription start site using the Eukaryotic Promoter Database [24]. We thus assessed HDAC1/7 binding to this SMAD binding site. Whereas no significant enrichment of HDAC1 was found, marked enrichment of HDAC7 was observed at putative SMAD binding sites relative to IgG control (Fig. 6I and S4B). These findings led us to examine the expression of HDAC7 in ccRCC in TCGA data. We observed that HDAC7 expression was significantly increased in ccRCC relative to normal kidney at the mRNA and protein levels (Fig. 6J and 6K). The increased protein expression of HDAC7 was validated in ccRCC as compared with patient-matched normal kidney samples (Fig. 6L). Collectively, these data support a novel role for HDAC7 in the silencing of TCA cycle enzymes in ccRCC.

**Figure 6.**
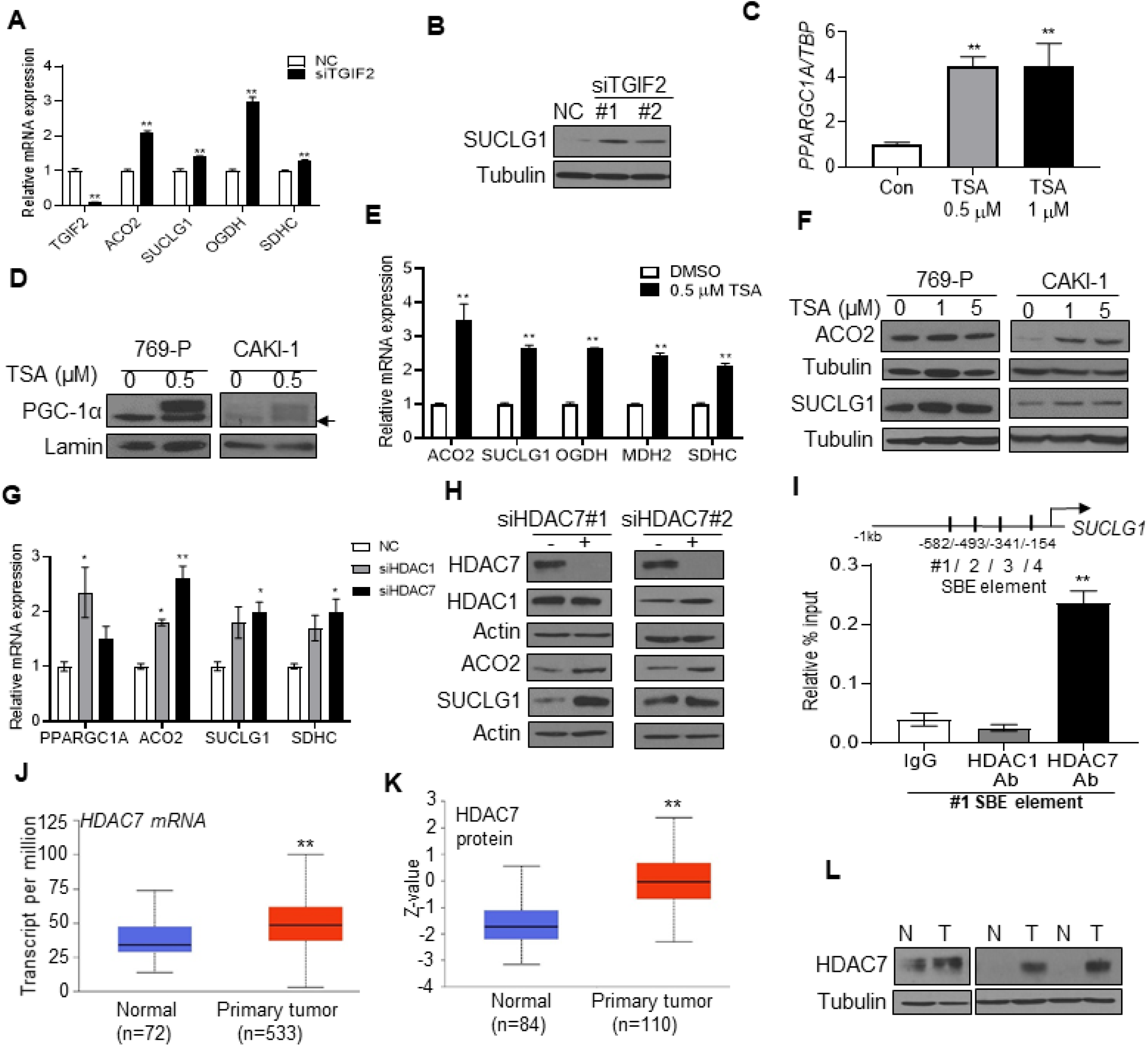
The expression of TCA cycle enzymes is restored by HDAC7 knockdown. **A and B)** The mRNA and protein expression of TCA cycle enzymes in CAKl-1 cells transfected with negative control (NC) or siRNA TGIF2 for 72 h. **C)** The mRNA expression of *PPARGC1A* in CAKl-1 cells treated with Trichostatin A (TSA) for 24 h (n=3/group). **D)** lmmunoblot analysis for PGC-1 a in nuclear lysate from 769-P and CAKl-1 cells treated with TSA for 24 h. Arrow represents non-specific band. **E)** The mRNA expression of TCA cycle enzymes in CAKl-1 cells treated with TSA for 24 h (n=3/group). **F)** lmmunoblot analysis for TCA cycle enzymes in cell lysate from 769-P and CAKl-1 cells treated with TSA for 24 h. **G)**769-P cells were transfected with indicated siRNA or negative control for 72 h. Relative mRNA expression was analyzed for TCA cycle enzymes (n=3/group). **H)** lmmunoblot analysis for the indicated proteins in cell lysates from 769-P cellstransfected with either NC or siRNA HDAC7 for 72 h. I) ChlP-qPCR was performed on CAKl-1 cells with mouse lgG, anti-HDAC1, and anti-HDAC7. The enriched DNA was quantified by qPCR with primer sets targeting the potential SMAD binding sites upstream of the *SUCLG1* transcription start site. Enrichment was calculated w th the percent input method (n=2/group, 3 independent experiments). J) RNA-Seq analysis of gene expression for *HDAC7*in the renal tumors in the TCGA data set. **K)** Protein expression of HDAC7 across pan-cancer subtypes from the CPTAC cohorts. **L)** lmmunoblot analysis of HDAC7 in patient-matched normal kidney (N) and tumor (T). All data represent 3 independent experiments and error bars are SEM. Asterisks indicate significant differences compared to control (\**P*<0.05,\*\**P*<0.01, One-way ANOVA with Tukey’s multiple comparisons test).

## DISCUSSION

The molecular basis by which tumor cells remodel their metabolism remains an area of intense investigation. Current understanding of metabolic remodeling in renal cancers has mainly focused on the HIF transcription factors [25]. ccRCC is associated with inactivation of the *VHL* gene due to genetic or epigenetic alterations. *VHL* inactivation results in stabilization of HIFs and subsequent upregulation of hypoxia-responsive genes that are involved in metabolic reprogramming including glycolytic enzymes. Although the role of *VHL* and *HIF* pathway is well described in RCC tumor initiation, inactivation of *VHL* alone may not be sufficient to drive ccRCC tumorigenesis [26]. Furthermore, prior studies demonstrate that HIF-1α expression is often lost in ccRCC and that it actually has tumor suppressive effects in the context of kidney cancer [25].

Despite the emphasis on increased expression of glycolytic enzymes in tumor metabolism, several lines of evidence support that alterations in mitochondrial metabolism are observed in renal cancer. For instance, prior studies have established decreased mitochondrial respiratory chain proteins in RCC and that respiratory complex chain activity is inversely correlated with prognosis [27, 28]. Loss of PGC-1α has been linked to these phenotypes [18]. In addition, germline loss of function mutations of the TCA cycle enzyme genes including *FH*, *SDHB*, *SDHC*, and *SDHD* have been identified in renal cancer that appear to have an increased risk for more aggressive tumors [29–31]. Notably, one of the major conclusions of the TCGA data analysis of ccRCC was that loss of TCA cycle enzyme expression is associated with poorer patient outcomes [32]. However, the precise mechanisms by which this occurs has not been reported. At the same time, the global suppression the TCA cycle indicates an epigenetic mechanism.

PGC-1α has been implicated in promoting tumor growth in breast, pancreas, and melanoma [33–35]. Alternatively, PGC-1α appear to suppress metastasis in prostate cancer and in a subset of melanomas [36, 37]. This inconsistency could be due to tissue-specific metabolic pathways that maintain tumor growth. We recently reported a novel role for PGC-1α in the suppression of collagen expression. Notably, several collagens are highly expressed in metastatic RCC [14]. Given that TGF-β signaling is known to promote collagen gene expression, we investigated the impact of TGF-β in RCC metabolism and tumor progression. TGF-β is a multifunctional extracellular cytokine that regulates cell growth, differentiation, migration and adhesion [38, 39]. TGF-β can inhibit cell proliferation and is considered tumor suppressive in early stages of tumorigenesis. In contrast, TGF-β can promote tumor progression through promoting epithelial-mesenchymal transition (EMT) [15, 40]. TGF-β signaling has been linked with metabolic reprogramming in cancer cells. For instance, increased expression or activity of glycolytic enzymes during TGF-β induced EMT has been demonstrated in multiple cancer cells [41–43]. Although TGF-β signaling stimulates glycolytic phenotypes during EMT process, the effects of TGF-β on the TCA cycle in cancer remains largely unknown. Therefore, our findings add novel insight into metabolic remodeling in RCC tumors. These are the first data, to our knowledge, to demonstrate an inhibitory effect of TGF-β on the gene expression of TCA cycle enzymes. Prior studies have implicated the impact of TGF-β on reduced *PPARGC1A* transcripts levels in renal fibrosis [44]. However, the detailed molecular mechanism of TGF-β mediated transcriptional regulation of *PPARGC1A* was not fully elucidated nor were the effects on the TCA cycle reported.

TGF-β signaling mainly regulates gene expression through receptor-mediated phosphorylation of SMAD proteins. Activated SMAD proteins translocate into the nucleus where they bind to DNA in the regulatory region of target genes. The affinity of SMADs for DNA is weak. Thus, SMADs often cooperates with additional transcription factors in the regulation of TGF-β responsive genes [45]. Our data implicate a novel role for TGIF2 in regulating the TCA cycle. Compelling evidence of the relevance of this axis is demonstrated by our *in vivo* studies which demonstrate that inhibition of TGF-β leads to increased TCA cycle enzyme expression in renal tumors. Our studies are among the first to demonstrate that mitochondrial aspects of the Warburg effect can be pharmacologically reversed *in vivo*.

Here, we also report that TGF-β works in concert with HDAC7 to suppress the expression of TCA cycle enzymes in RCC. These are the first data to reveal that HDAC7 can suppress genes that are involved in mitochondrial metabolism. Although we found the increased mRNA expression of *PPARGC1A* in either HDAC1 or HDAC7 knockdown, no significant enrichment of these HDACs in the promoter region of *PPARGC1A* was found (data not shown). Thus, our findings support that HDAC7 mediates its effects directly at TCA cycle enzyme genes. These data indicate an alternate means by which mitochondrial metabolism could be activated in RCC. As supportive evidence, recent studies in glioma cells indicate that HDAC inhibitors can activate mitochondrial metabolism [46]. Future studies will focus on the role mitochondrial metabolism plays in tumor biology as well as response to therapies including immune checkpoint blockade.

In summary, our findings provide a novel insight into the molecular mechanisms driving metabolic reprogramming in renal cancer. Our studies demonstrate that TGF-β signaling represses *PPARGC1A* and TCA cycle enzymes and are the first to demonstrate a role for TGIF2/HDAC7 in the suppression of mitochondrial metabolism. Hence, these findings highlight an intriguing possibility that this axis could be targeted by epigenetic based therapies which are currently in use or in clinical trials.

## MATERIALS AND METHODS

### Cell culture

RCC cell lines (RCC10, CAKI-1, 769-P, and 786-0) were purchased from ATCC except for RCC4 (kindly provided by P. Ratcliffe, University of Oxford), and RXF-393 (NCI). All RCC cell lines were maintained as described previously [14]. HK2 renal epithelial cells were purchased from ATCC. Primary renal proximal tubule epithelial cells (RPTEC) were acquired from Lonza and grown in renal epithelial cell growth basal medium supplemented with 10% fetal bovine serum and penicillin streptomycin (100 U/mL) in 5% CO_2_ at 37°C. Cells were used within 10 passages of the initial stock and periodically screened for mycoplasma contamination.

### TGF-β inhibitor treatment

RCC cells were grown in serum-free medium supplemented with 0.1% bovine serum albumin (Roche) and 2 mM glutamine for 24 h prior to treatment with either TGF-β (1 ng/mL, Millipore Calbiochem) or 10 uM TGF-β inhibitors (SB431542; Sigma-Aldrich, LY2109761 and LY364947; Selleckchem) for the indicated times.

### siRNA transfection

RCC cells were transfected with 25 nM of a negative control siRNA or siRNA indicated target genes using Lipofectamine RNAiMAX reagent (Invitrogen) for 72 h.

### Plasmid and virus infections

Human *PPARGC1A* cDNA was obtained from GeneCopoeia. Lentiviral shRNA constructs for *PPARGC1A* were purchased from Sigma. Lentiviral particles were generated by co-transfecting HEK293T cells with packaging plasmids using the calcium phosphate method. The detailed methods were described previously [14]. For adenoviral studies, RCC cells were transduced with either GFP or *PPARGC1A* (Vector Biolabs).

### Cellular Bioenergetics

Cellular bioenergetics was determined using the Seahorse XFe96 Analyzer (Agilent Technologies). RCC cells were treated with DMSO or TGF-β inhibitors for 24 h prior to being plated in a Seahorse XF96 plate (25 × 10^3^ cells per well; n=8/group). Cells rested overnight before measuring the oxygen consumption rate (OCR) the following day. Cells were then washed with extracellular flux media and allowed to equilibrate for 1 h prior to being exposed to the mitochondrial stress test.

### Patient samples and gene expression profiling

Patient samples for gene expression profiling (normal=9, primary=9, metastasis n=26) were not patient-matched and are previously described [12]. A separate cohort of patient-matched samples (n=6/group) for the validation of gene expression profiling was obtained from UAB Hospital and has been described previously [12]. All studies were performed in accordance with the institutional IRB.

### Quantitative RT-PCR

Total RNA from human and mouse kidney samples was harvested using the RNeasy Mini Kit (Qiagen). Total RNA isolation from cultured cells was extracted using Trizol reagent (Ambion) according to the manufacturer’s instructions. cDNA was generated using a High-Capacity cDNA Reverse Transcription Kit (Applied Biosystems). Quantitative RT-PCR was performed using the indicated Taqman primers in QuantStudio 6K Flex Real-Time PCR System (Applied Biosystem) (Table S1A). The mRNA expression of target gene was normalized to either ribosomal protein (*RPLPO*) or TATA binding protein (*TBP*). The normalized Ct value was quantified using the double delta Ct analysis.

### ChIP-qPCR

The potential SMAD binding sites near the *SUCLG1* transcriptional start site were identified using the Eukaryotic Promoter Database [24]. The ChIP experiment was conducted using the EZ-Magna ChIP Chromatin IP A/G kit (Millipore) according to the manufacturer’s instructions. After pull-down with either mouse IgG or HDAC1/7 antibody, input DNA and immunoprecipitated DNA were purified (Zymo Research). The target DNA enrichment was calculated based on the % input method. Primer sequences for ChIP assay are described in Table S1B.

### Measurement of mitochondrial DNA content

DNA from RCC cells was extracted with the QIAamp DNA mini kit (Qiagene). Mitochondrial DNA content was analyzed by measuring the relative levels of mitochondrial DNA D-Loop and mitochondrial DNA-encoded *MT-CO2* by qRT-PCR. Genomic DNA-encoded β-actin was used as a normalizer. The list of primers used for this study are presented in Table S2A.

### Immunoblotting analysis

RCC cells were lysed with ice-cold RIPA buffer containing 1X protease inhibitor (Halt protease and phosphatase inhibitor cocktail, ThermoFisher). Human and mouse kidney samples were homogenized with microbeads (Bioexpress) containing SDS lysis buffer. Preparation of samples and Western blot analysis were described previously [12, 14]. Antibodies used in this study are described in Table S2B.

### 13C glucose incorporation analysis

CAKI-1 cells were grown to approximately 80% confluence. Cells were supplemented with fresh 5.5 mM [U-13C6] glucose (Cambridge Isotope Laboratories). After 24 hr incubation, metabolites were extracted using cold 80% HPLC graded methanol. The sample was centrifuged at 20,000g for 10 min and the supernatant was dried under vacuum. Pellets were reconstituted in solvent (water:methanol:acetonitrile, 2:1:1, v/v) and further analyzed by LC-MS as previously described [14, 47].

### Orthotopic tumor challenge in vivo

Orthotopic tumor challenges were performed using 5-week old SCID male mice (Charles River Laboratories). Luciferase-expressing CAKI-1 cells (1.5×10^6^) were mixed in a 1:1 ratio with Matrigel and injected under the left renal capsule. Bioluminescent imaging was performed after one week to assess for xenograft formation. For TGF-β inhibition *in vivo*, SB-431542 (Sigma-Aldrich) was dissolved with 20% DMSO and filtered through a 0.45 μm filter (Milipore). Mice received either 20% DMSO as a control group or diluted SB-431542 in 20% DMSO (10 mg/kg) three times per week for 5 weeks (n=3/group). Tumor progression was weekly evaluated by measuring the luciferase signal with IVIS Lumina III In Vivo System (PerkinElmer). All animal studies were conducted in accordance with the NIH guidelines and were approved by UAB Institutional Animal Care and Use Committee (IACUC).

## ACKNOWLEDGMENTS

The research reported in this article was supported by Department of Veteran Affairs grant BX002930 and NIH/NCI grant R01CA20053. This work was also supported by the NIH (P30 CA013148).

## REFERENCES

1 Janzen, N.K., et al., Surveillance after radical or partial nephrectomy for localized renal cell carcinoma and management of recurrent disease. Urol Clin North Am, 2003. 30(4): p. 843–52.

2 Jemal, A., et al., Cancer statistics, 2005. CA Cancer J Clin, 2005. 55(1): p. 10–30.

3 Kibel, A., et al., Binding of the von Hippel-Lindau tumor suppressor protein to Elongin B and C. Science, 1995. 269(5229): p. 1444–6.

4 Ivan, M., et al., HIFalpha targeted for VHL-mediated destruction by proline hydroxylation: implications for O2 sensing. Science, 2001. 292(5516): p. 464–8.

5 Kaelin, W.G., Jr., The von Hippel-Lindau protein, HIF hydroxylation, and oxygen sensing. Biochem Biophys Res Commun, 2005. 338(1): p. 627–38.

6 Semenza, G.L., et al., Transcriptional regulation of genes encoding glycolytic enzymes by hypoxia-inducible factor 1. J Biol Chem, 1994. 269(38): p. 23757–63.

7 Denko, N.C., Hypoxia, HIF1 and glucose metabolism in the solid tumour. Nat Rev Cancer, 2008. 8(9): p. 705–13.

8 Hu, J., et al., Heterogeneity of tumor-induced gene expression changes in the human metabolic network. Nat Biotechnol, 2013. 31(6): p. 522–9.

9 Sudarshan, S., et al., Fumarate hydratase deficiency in renal cancer induces glycolytic addiction and hypoxia-inducible transcription factor 1alpha stabilization by glucose-dependent generation of reactive oxygen species. Mol Cell Biol, 2009. 29(15): p. 4080–90.

10 Courtney, K.D., et al., Isotope Tracing of Human Clear Cell Renal Cell Carcinomas Demonstrates Suppressed Glucose Oxidation In Vivo. Cell Metab, 2018. 28(5): p. 793–800 e2.

11 Comprehensive molecular characterization of clear cell renal cell carcinoma. Nature, 2013. 499(7456): p. 43–9.

12 Nam, H.Y., et al., Integrative Epigenetic and Gene Expression Analysis of Renal Tumor Progression to Metastasis. Mol Cancer Res, 2019. 17(1): p. 84–96.

13 Puigserver, P., et al., A cold-inducible coactivator of nuclear receptors linked to adaptive thermogenesis. Cell, 1998. 92(6): p. 829–39.

14 Nam, H., et al., PGC1alpha suppresses kidney cancer progression by inhibiting collagen-induced SNAIL expression. Matrix Biol, 2020.

15 Massague, J., TGFbeta in Cancer. Cell, 2008. 134(2): p. 215–30.

16 Chen, F., et al., Pan-cancer molecular subtypes revealed by mass-spectrometry-based proteomic characterization of more than 500 human cancers. Nat Commun, 2019. 10(1): p. 5679.

17 Papandreou, I., et al., HIF-1 mediates adaptation to hypoxia by actively downregulating mitochondrial oxygen consumption. Cell Metab, 2006. 3(3): p. 187–97.

18 LaGory, E.L., et al., Suppression of PGC-1alpha Is Critical for Reprogramming Oxidative Metabolism in Renal Cell Carcinoma. Cell Rep, 2015. 12(1): p. 116–127.

19 Akiyoshi, S., et al., c-Ski acts as a transcriptional co-repressor in transforming growth factor-beta signaling through interaction with smads. J Biol Chem, 1999. 274(49): p. 35269–77.

20 Luo, K., et al., The Ski oncoprotein interacts with the Smad proteins to repress TGFbeta signaling. Genes Dev, 1999. 13(17): p. 2196–206.

21 Wotton, D., et al., A Smad transcriptional corepressor. Cell, 1999. 97(1): p. 29–39.

22 Cai, L., et al., Genomic regression analysis of coordinated expression. Nat Commun, 2017. 8(1): p. 2187.

23 Melhuish, T.A., C.M. Gallo, and D. Wotton, TGIF2 interacts with histone deacetylase 1 and represses transcription. J Biol Chem, 2001. 276(34): p. 32109–14.

24 Dreos, R., et al., The eukaryotic promoter database in its 30th year: focus on non-vertebrate organisms. Nucleic Acids Res, 2017. 45(D1): p. D51–D55.

25 Shen, C., et al., Genetic and functional studies implicate HIF1alpha as a 14q kidney cancer suppressor gene. Cancer Discov, 2011. 1(3): p. 222–35.

26 Nickerson, M.L., et al., Improved identification of von Hippel-Lindau gene alterations in clear cell renal tumors. Clin Cancer Res, 2008. 14(15): p. 4726–34.

27 Meierhofer, D., et al., Decrease of mitochondrial DNA content and energy metabolism in renal cell carcinoma. Carcinogenesis, 2004. 25(6): p. 1005–10.

28 Simonnet, H., et al., Low mitochondrial respiratory chain content correlates with tumor aggressiveness in renal cell carcinoma. Carcinogenesis, 2002. 23(5): p. 759–68.

29 Grubb, R.L., 3rd, et al., Hereditary leiomyomatosis and renal cell cancer: a syndrome associated with an aggressive form of inherited renal cancer. J Urol, 2007. 177(6): p. 2074–9; discussion 2079-80.

30 Ricketts, C.J., et al., Succinate dehydrogenase kidney cancer: an aggressive example of the Warburg effect in cancer. J Urol, 2012. 188(6): p. 2063–71.

31 Tomlinson, I.P., et al., Germline mutations in FH predispose to dominantly inherited uterine fibroids, skin leiomyomata and papillary renal cell cancer. Nat Genet, 2002. 30(4): p. 406–10.

32 Cancer Genome Atlas Research, N., Comprehensive molecular characterization of clear cell renal cell carcinoma. Nature, 2013. 499(7456): p. 43–9.

33 LeBleu, V.S., et al., PGC-1alpha mediates mitochondrial biogenesis and oxidative phosphorylation in cancer cells to promote metastasis. Nat Cell Biol, 2014. 16(10): p. 992–1003, 1-15.

34 Sancho, P., et al., MYC/PGC-1alpha Balance Determines the Metabolic Phenotype and Plasticity of Pancreatic Cancer Stem Cells. Cell Metab, 2015. 22(4): p. 590–605.

35 Haq, R., et al., Oncogenic BRAF regulates oxidative metabolism via PGC1alpha and MITF. Cancer Cell, 2013. 23(3): p. 302–15.

36 Luo, C., et al., A PGC1alpha-mediated transcriptional axis suppresses melanoma metastasis. Nature, 2016. 537(7620): p. 422–426.

37 Torrano, V., et al., The metabolic co-regulator PGC1alpha suppresses prostate cancer metastasis. Nat Cell Biol, 2016. 18(6): p. 645–656.

38 Dongre, A. and R.A. Weinberg, New insights into the mechanisms of epithelial-mesenchymal transition and implications for cancer. Nat Rev Mol Cell Biol, 2019. 20(2): p. 69–84.

39 Nieto, M.A., et al., Emt: 2016. Cell, 2016. 166(1): p. 21–45.

40 Ikushima, H. and K. Miyazono, TGFbeta signalling: a complex web in cancer progression. Nat Rev Cancer, 2010. 10(6): p. 415–24.

41 Li, W., et al., Increased 18F-FDG uptake and expression of Glut1 in the EMT transformed breast cancer cells induced by TGF-beta. Neoplasma, 2010. 57(3): p. 234–40.

42 Liu, M., et al., Epithelial-mesenchymal transition induction is associated with augmented glucose uptake and lactate production in pancreatic ductal adenocarcinoma. Cancer Metab, 2016. 4: p. 19.

43 Rodriguez-Garcia, A., et al., TGF-beta1 targets Smad, p38 MAPK, and PI3K/Akt signaling pathways to induce PFKFB3 gene expression and glycolysis in glioblastoma cells. FEBS J, 2017. 284(20): p. 3437–3454.

44 Kang, H.M., et al., Defective fatty acid oxidation in renal tubular epithelial cells has a key role in kidney fibrosis development. Nat Med, 2015. 21(1): p. 37–46.

45 Shi, Y., et al., Crystal structure of a Smad MH1 domain bound to DNA: insights on DNA binding in TGF-beta signaling. Cell, 1998. 94(5): p. 585–94.

46 Nguyen, T.T.T., et al., HDAC inhibitors elicit metabolic reprogramming by targeting super-enhancers in glioblastoma models. J Clin Invest, 2020. 130(7): p. 3699–3716.

47 Brinkley, G., et al., Teleological role of L-2-hydroxyglutarate dehydrogenase in the kidney. Dis Model Mech, 2020. 13(11).

