## Supplemental files for "TGF-β signaling suppresses TCA cycle metabolism in renal cancer"

### Slide 1
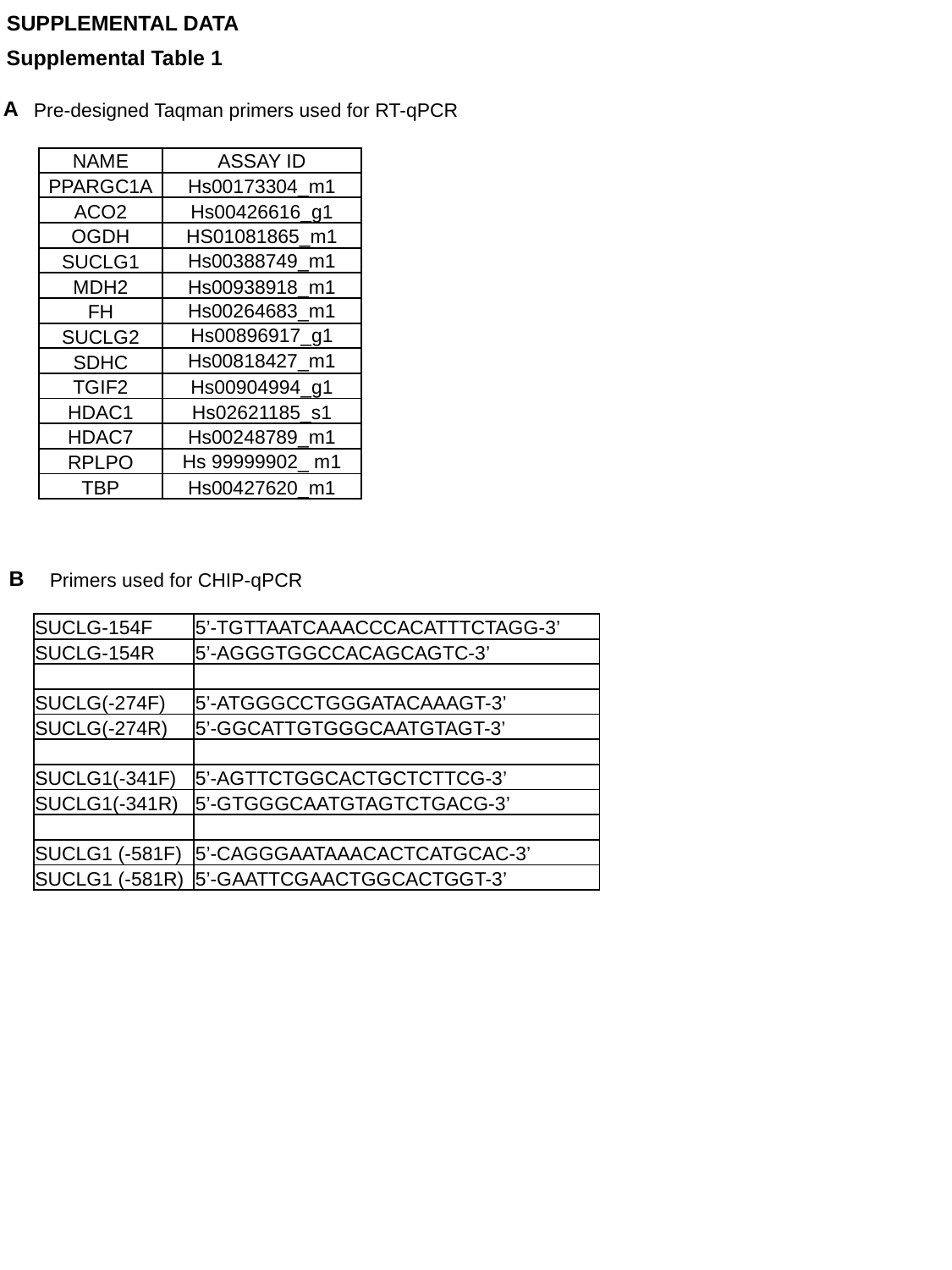

SUPPLEMENTAL DATA
Supplemental Table 1
A
Pre-designed Taqman primers used for RT-qPCR
| NAME | ASSAY ID |
| --- | --- |
| PPARGC1A | Hs00173304\_m1 |
| ACO2 | Hs00426616\_g1 |
| OGDH | HS01081865\_m1 |
| SUCLG1 | Hs00388749\_m1 |
| MDH2 | Hs00938918\_m1 |
| FH | Hs00264683\_m1 |
| SUCLG2 | Hs00896917\_g1 |
| SDHC | Hs00818427\_m1 |
| TGIF2 | Hs00904994\_g1 |
| HDAC1 | Hs02621185\_s1 |
| HDAC7 | Hs00248789\_m1 |
| RPLPO | Hs 99999902\_ m1 |
| TBP | Hs00427620\_m1 |
B
Primers used for CHIP-qPCR
| SUCLG-154F | 5’-TGTTAATCAAACCCACATTTCTAGG-3’ |
| --- | --- |
| SUCLG-154R | 5’-AGGGTGGCCACAGCAGTC-3’ |
| SUCLG(-274F) | 5’-ATGGGCCTGGGATACAAAGT-3’ |
| SUCLG(-274R) | 5’-GGCATTGTGGGCAATGTAGT-3’ |
| SUCLG1(-341F) | 5’-AGTTCTGGCACTGCTCTTCG-3’ |
| SUCLG1(-341R) | 5’-GTGGGCAATGTAGTCTGACG-3’ |
| SUCLG1 (-581F) | 5’-CAGGGAATAAACACTCATGCAC-3’ |
| SUCLG1 (-581R) | 5’-GAATTCGAACTGGCACTGGT-3’ |

### Slide 2
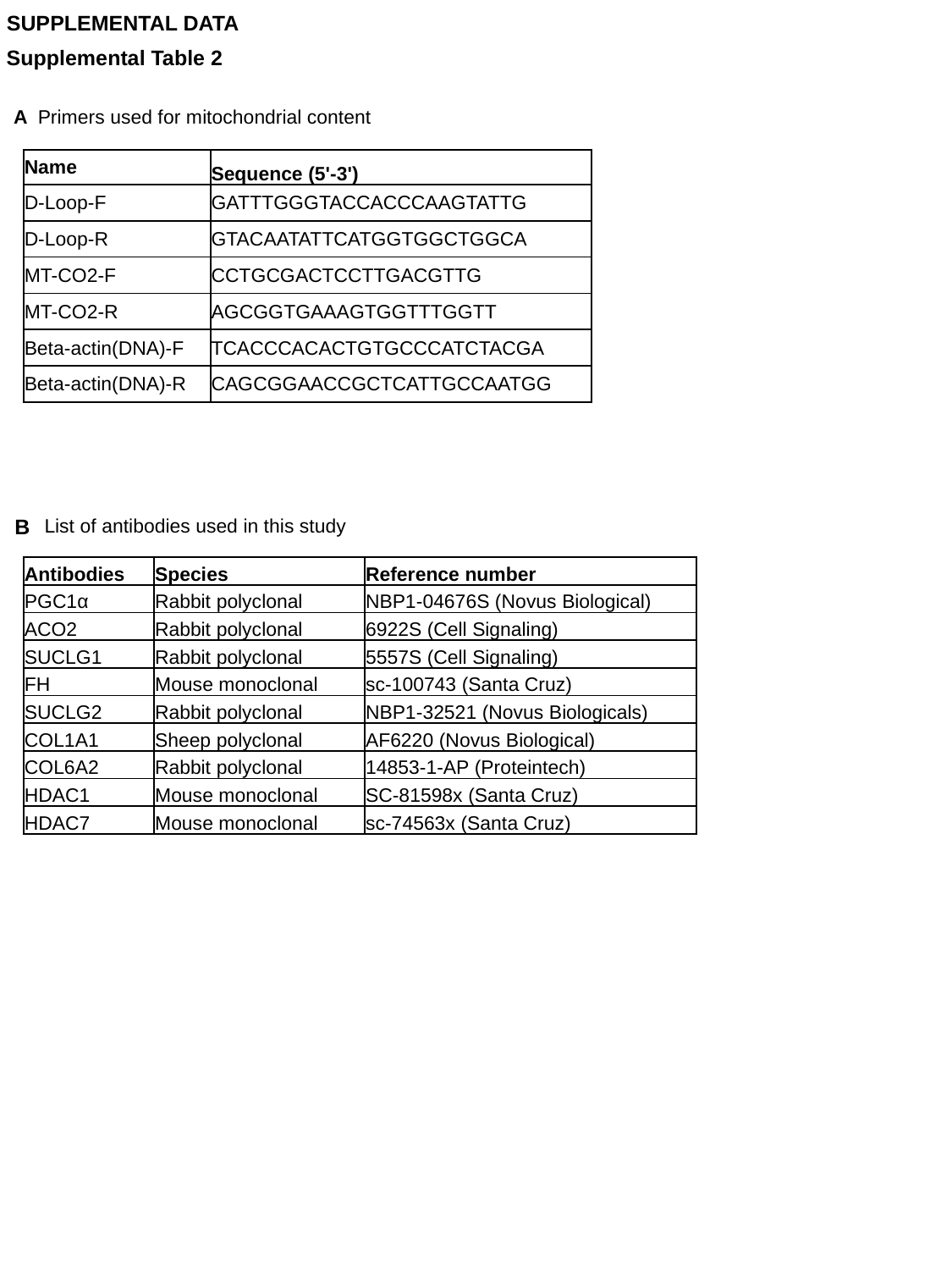

SUPPLEMENTAL DATA
Supplemental Table 2
A Primers used for mitochondrial content
| Name | Sequence (5'-3') |
| --- | --- |
| D-Loop-F | GATTTGGGTACCACCCAAGTATTG |
| D-Loop-R | GTACAATATTCATGGTGGCTGGCA |
| MT-CO2-F | CCTGCGACTCCTTGACGTTG |
| MT-CO2-R | AGCGGTGAAAGTGGTTTGGTT |
| Beta-actin(DNA)-F | TCACCCACACTGTGCCCATCTACGA |
| Beta-actin(DNA)-R | CAGCGGAACCGCTCATTGCCAATGG |
B
List of antibodies used in this study
| Antibodies | Species | Reference number |
| --- | --- | --- |
| PGC1α | Rabbit polyclonal | NBP1-04676S (Novus Biological) |
| ACO2 | Rabbit polyclonal | 6922S (Cell Signaling) |
| SUCLG1 | Rabbit polyclonal | 5557S (Cell Signaling) |
| FH | Mouse monoclonal | sc-100743 (Santa Cruz) |
| SUCLG2 | Rabbit polyclonal | NBP1-32521 (Novus Biologicals) |
| COL1A1 | Sheep polyclonal | AF6220 (Novus Biological) |
| COL6A2 | Rabbit polyclonal | 14853-1-AP (Proteintech) |
| HDAC1 | Mouse monoclonal | SC-81598x (Santa Cruz) |
| HDAC7 | Mouse monoclonal | sc-74563x (Santa Cruz) |

### Slide 3
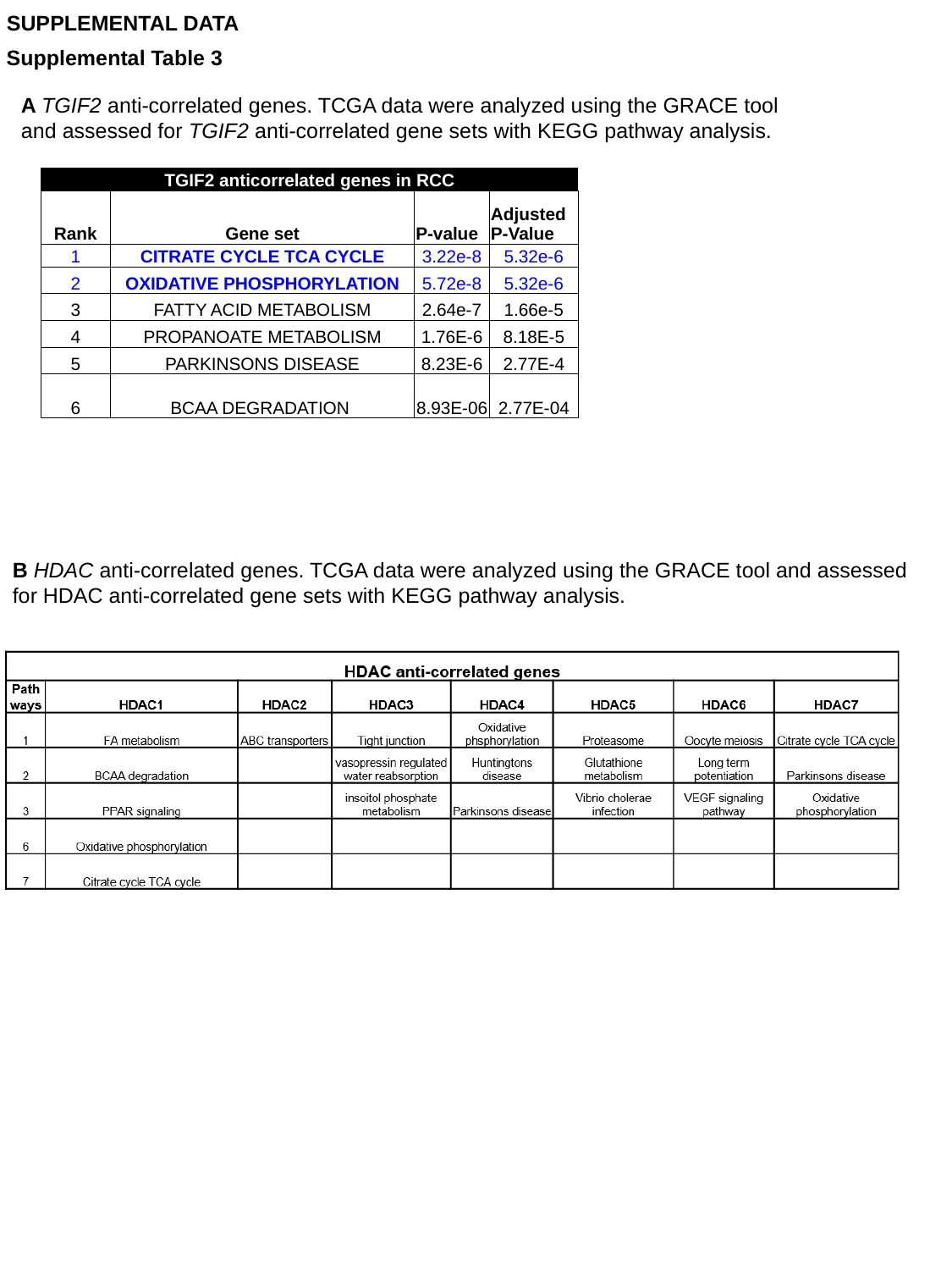

SUPPLEMENTAL DATA
Supplemental Table 3
A TGIF2 anti-correlated genes. TCGA data were analyzed using the GRACE tool
and assessed for TGIF2 anti-correlated gene sets with KEGG pathway analysis.
| TGIF2 anticorrelated genes in RCC | | | |
| --- | --- | --- | --- |
| Rank | Gene set | P-value | Adjusted P-Value |
| 1 | CITRATE CYCLE TCA CYCLE | 3.22e-8 | 5.32e-6 |
| 2 | OXIDATIVE PHOSPHORYLATION | 5.72e-8 | 5.32e-6 |
| 3 | FATTY ACID METABOLISM | 2.64e-7 | 1.66e-5 |
| 4 | PROPANOATE METABOLISM | 1.76E-6 | 8.18E-5 |
| 5 | PARKINSONS DISEASE | 8.23E-6 | 2.77E-4 |
| 6 | BCAA DEGRADATION | 8.93E-06 | 2.77E-04 |
B HDAC anti-correlated genes. TCGA data were analyzed using the GRACE tool and assessed for HDAC anti-correlated gene sets with KEGG pathway analysis.

### Slide 4
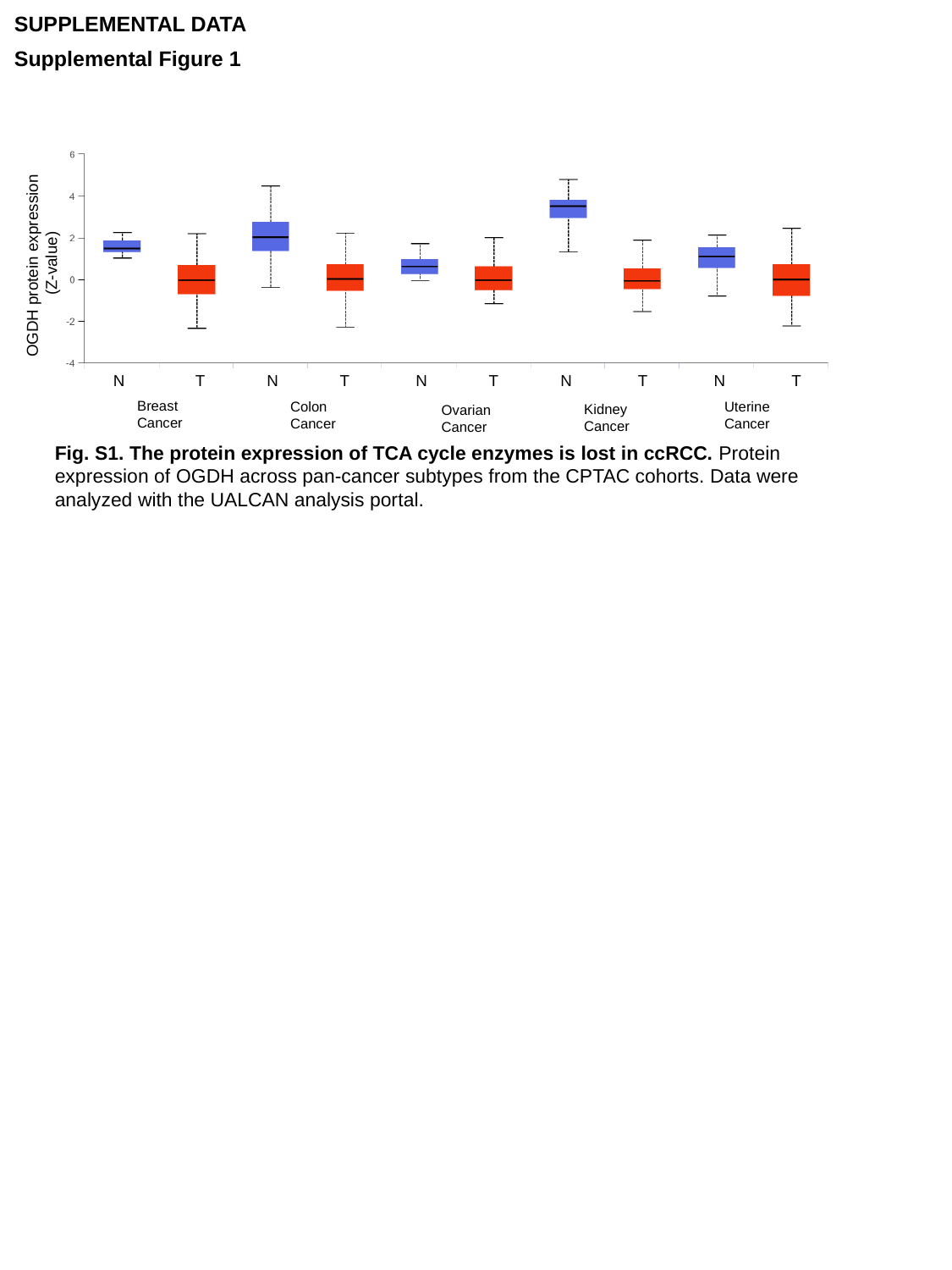

SUPPLEMENTAL DATA
Supplemental Figure 1
OGDH protein expression
(Z-value)
 N T N T N T N T N T
Breast
Cancer
Colon
Cancer
Uterine
Cancer
Kidney
Cancer
Ovarian
Cancer
Fig. S1. The protein expression of TCA cycle enzymes is lost in ccRCC. Protein expression of OGDH across pan-cancer subtypes from the CPTAC cohorts. Data were analyzed with the UALCAN analysis portal.

### Slide 5
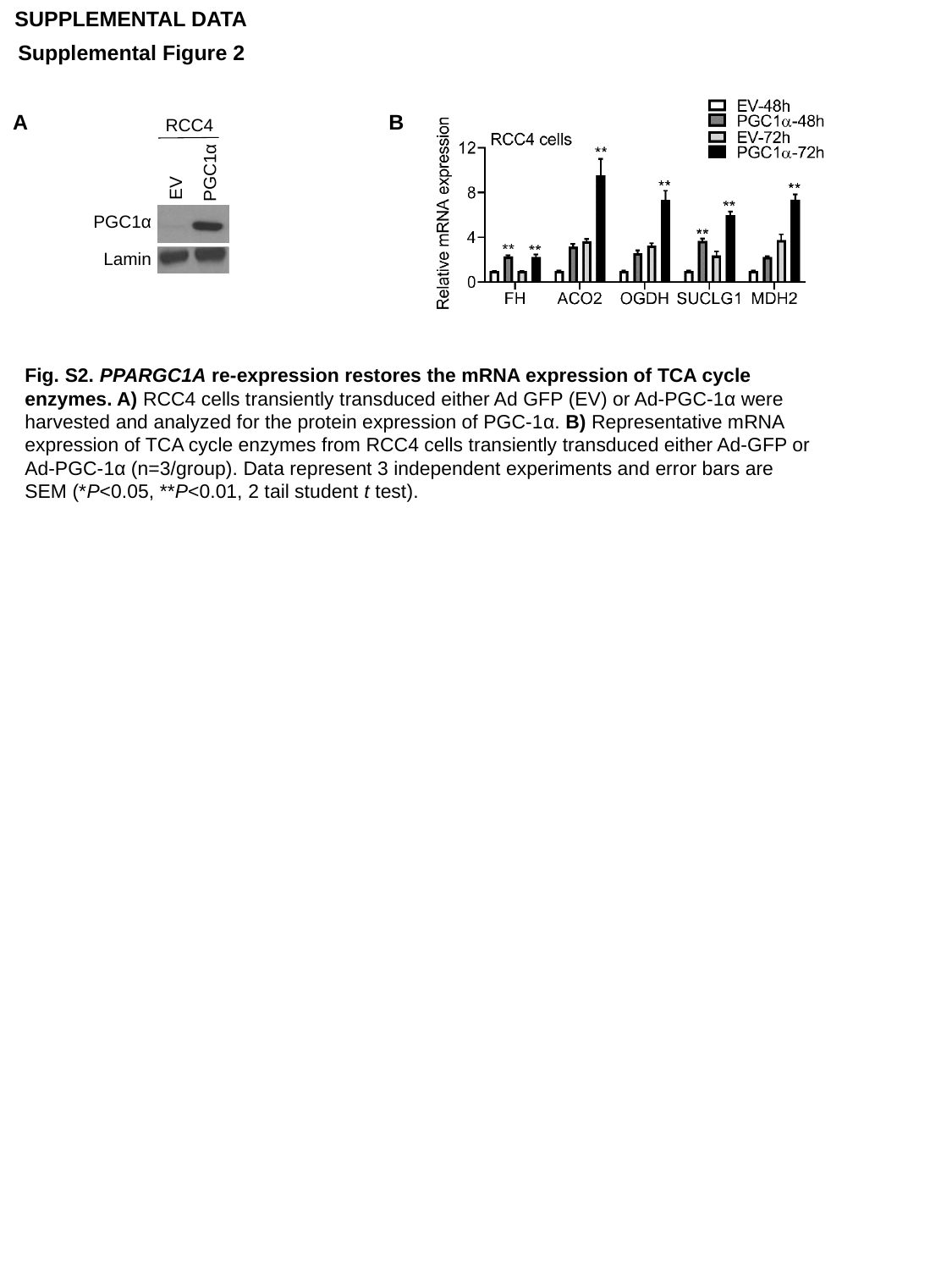

SUPPLEMENTAL DATA
Supplemental Figure 2
B
A
RCC4
PGC1α
EV
PGC1α
Lamin
Fig. S2. PPARGC1A re-expression restores the mRNA expression of TCA cycle enzymes. A) RCC4 cells transiently transduced either Ad GFP (EV) or Ad-PGC-1α were harvested and analyzed for the protein expression of PGC-1α. B) Representative mRNA expression of TCA cycle enzymes from RCC4 cells transiently transduced either Ad-GFP or Ad-PGC-1α (n=3/group). Data represent 3 independent experiments and error bars are SEM (*P<0.05, **P<0.01, 2 tail student t test).

### Slide 6
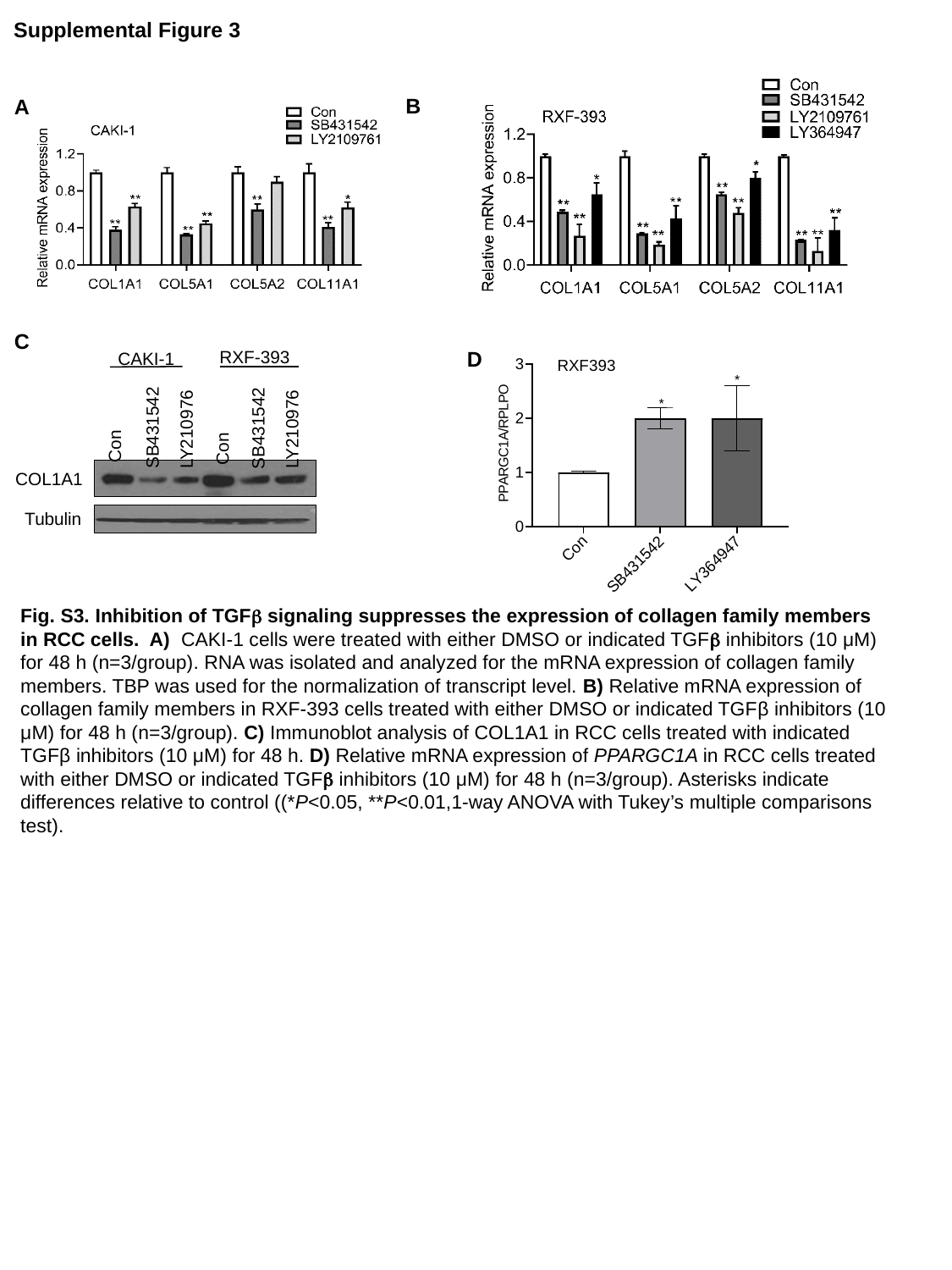

Supplemental Figure 3
B
A
C
RXF-393
CAKI-1
SB431542
LY210976
Con
SB431542
LY210976
Con
COL1A1
Tubulin
D
RXF393
Fig. S3. Inhibition of TGFβ signaling suppresses the expression of collagen family members in RCC cells. A) CAKI-1 cells were treated with either DMSO or indicated TGFβ inhibitors (10 μM) for 48 h (n=3/group). RNA was isolated and analyzed for the mRNA expression of collagen family members. TBP was used for the normalization of transcript level. B) Relative mRNA expression of collagen family members in RXF-393 cells treated with either DMSO or indicated TGFβ inhibitors (10 μM) for 48 h (n=3/group). C) Immunoblot analysis of COL1A1 in RCC cells treated with indicated TGFβ inhibitors (10 μM) for 48 h. D) Relative mRNA expression of PPARGC1A in RCC cells treated with either DMSO or indicated TGFβ inhibitors (10 μM) for 48 h (n=3/group). Asterisks indicate differences relative to control ((*P<0.05, **P<0.01,1-way ANOVA with Tukey’s multiple comparisons test).

### Slide 7
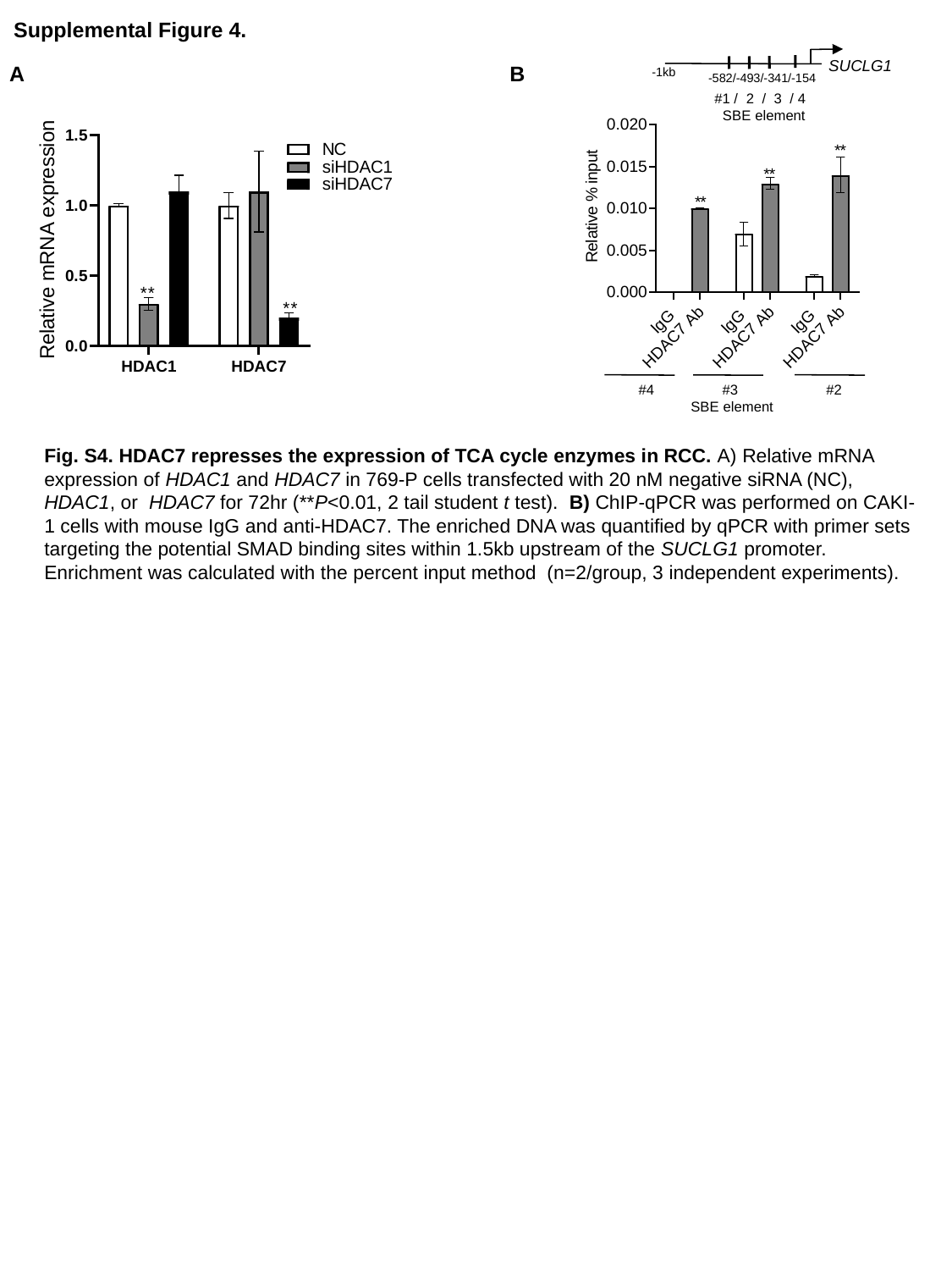

Supplemental Figure 4.
-1kb
-582/-493/-341/-154
SUCLG1
#1 / 2 / 3 / 4
 SBE element
B
A
 #4 #3 #2
 SBE element
Fig. S4. HDAC7 represses the expression of TCA cycle enzymes in RCC. A) Relative mRNA expression of HDAC1 and HDAC7 in 769-P cells transfected with 20 nM negative siRNA (NC), HDAC1, or HDAC7 for 72hr (**P<0.01, 2 tail student t test). B) ChIP-qPCR was performed on CAKI-1 cells with mouse IgG and anti-HDAC7. The enriched DNA was quantified by qPCR with primer sets targeting the potential SMAD binding sites within 1.5kb upstream of the SUCLG1 promoter. Enrichment was calculated with the percent input method (n=2/group, 3 independent experiments).
